# A GO catalogue of human DNA-binding transcription factors

**DOI:** 10.1101/2020.10.28.359232

**Authors:** Ruth C. Lovering, Pascale Gaudet, Marcio L. Acencio, Alex Ignatchenko, Arttu Jolma, Oriol Fornes, Martin Kuiper, Ivan V. Kulakovskiy, Astrid Lægreid, Maria J. Martin, Colin Logie

**Affiliations:** Functional Gene Annotation, Preclinical and Fundamental Science, UCL Institute of Cardiovascular Science, University College London, London, UK; Swiss-Prot group, SIB Swiss Institute of Bioinformatics, 1 Rue Michel-Servet, 1211 Geneve, Switzerland; Department of Clinical and Molecular Medicine, Norwegian University of Science and Technology, Trondheim, Norway; Current affiliation: Bioinformatics Core Unit, Luxembourg Centre for Systems Biomedicine, Université du Luxembourg, Esch-sur-Alzette, Luxembourg; European Molecular Biology Laboratory, European Bioinformatics Institute (EMBL-EBI), Wellcome Genome Campus, Hinxton, Cambridge CB10 1SD, UK; Donnelly Centre, University of Toronto, Toronto, ON, Canada; Centre for Molecular Medicine and Therapeutics, Department of Medical Genetics, BC Children’s Hospital Research Institute, University of British Columbia, 950 W 28th Ave, Vancouver, BC V5Z 4H4, Canada; Department of Biology, Norwegian University of Science and Technology, Trondheim, Norway; Institute of Protein Research, Russian Academy of Sciences, Institutskaya 4, Pushchino, 142290, Russia; Engelhardt Institute of Molecular Biology, Russian Academy of Sciences, Vavilova 32, Moscow, GSP-1, 119991, Russia; Molecular Biology Department, Faculty of Science, Radboud University, PO box 9101, 6500HB Nijmegen, The Netherlands.

**Keywords:** Biocuration, Gene Ontology, DNA-binding transcription factor

## Abstract

DNA-binding transcription factors recognise genomic addresses, specific sequence motifs in gene regulatory regions, to control gene transcription. A complete and reliable catalogue of all DNA-binding transcription factors is key to investigating the delicate balance of gene regulation in response to environmental and developmental stimuli. The need for such a catalogue of proteins is demonstrated by the many lists of DNA-binding transcription factors that have been produced over the past decade.

The COST Action Gene Regulation Ensemble Effort for the Knowledge Commons (GREEKC) Consortium brought together experts in the field of transcription with the aim of providing high quality and interoperable gene regulatory data. The Gene Ontology (GO) Consortium provides strict definitions for gene product function, including factors that regulate transcription. The collaboration between the GREEKC and GO Consortia has enabled the application of those definitions to produce a new curated catalogue of human DNA-binding transcription factors, that can be accessed at https://www.ebi.ac.uk/QuickGO/targetset/dbTF.

In addition, this curation effort has led to the GO annotation of almost sixty thousand DNA-binding transcription factors in over a hundred species. Thus, this work will aid researchers investigating the regulation of transcription in both biomedical and basic science.

## Introduction

The expression of house-keeping genes, as well as the regulated expression of gene products in response to environmental and developmental conditions, is controlled by carefully tuned cellular events. While signal transduction, mRNA splicing, translation, post-translational processing, and targeting to the appropriate cellular location are all potential points for regulation, the control of transcription is a master switch. Ultimately, the complement of genes expressed in a cell determines its identity and can also indicate or cause diseases. It is now recognised that exon variants contribute far less to inherited diseases or disease risk than dysregulation of gene expression (1,2). Consequently, non-coding genomic variants associated with disease are now of major interest in biomedical research. Many of those non-coding variants lie in transcription regulatory regions, hence the proteins and RNAs that bind these regions may serve as promising novel drug targets (3–5). Identifying all sequence-specific DNA-binding transcription factors (dbTFs) will provide the foundations for understanding the complex processes involved in ensuring properly regulated gene expression and recognizing conditions under which aberrant gene regulation can be treated therapeutically.

The Gene Ontology (GO) Consortium resource is widely used for functional analysis of high-throughput datasets. GO groups genes according to three biological aspects: the molecular activity of a gene product (GO Molecular Function), its role in the cell or whole organism (GO Biological Process), and the location of its activity in the cellular environment (GO Cellular Component) (6,7). Members of the GO Consortium have established various methods to associate gene products with GO terms. All of these approaches use manual curation to some extent (8). While some annotations result from the extraction of knowledge from individual published articles by biocurators, the majority of GO annotations are created by automatic pipelines. These pipelines either map experimentally-derived annotations to orthologous genes in different species (9), or apply sets of GO terms to all gene products which contain well-characterised protein domains (10). Finally, a significant number of annotations are produced by phylogenetic inference (11).

GO defines several key activities required for transcription and its regulation. General transcription factors (GTFs) provide the constitutive machinery required for transcription to occur, whereas, sequence-specific DNA-binding transcription factors (dbTFs) and transcription coregulators (coTFs) function to limit or increase gene expression. These activities converge on the RNA polymerase transcription cycle, from factors that determine the accessibility of the chromatin regions associated with the gene, to assembly of its pre-initiation complexes and to the elongating enzyme complexes (12). In GO, all of the above molecular activity terms have more specific descendants: for example, the dbTF parent term ‘DNA-binding transcription factor activity’ (GO:0003700) has a more specific term for eukaryotic protein-coding genes: ‘DNA-binding transcription factor activity, RNA polymerase II-specific’ (GO:0000981) (Gaudet et al., 2020a, in preparation).

Experts agree that there are approximately 1,500 dbTFs in the human genome (13,14). However, at the beginning of this work it was clear that the GO annotation dataset had not captured the dbTF activity of many of these proteins, as only around 1,300 human proteins were associated with a dbTF GO term. In addition, there was a concern that some proteins were inappropriately associated with a dbTF term. As the quality and coverage of annotations directly impact the interpretation of high-throughput data analysis (15), it was necessary to address these inconsistencies. There are several reasons why consistent association of a dbTF activity GO term with sequence-specific DNA-binding transcription factors has proved to be particularly challenging. First, many papers describe the characterised protein as a transcription factor, without specifying whether the activity of the protein involves direct DNA contacts. Second, many experiments used to investigate dbTFs rely on existing knowledge, which is not always conveyed in the article that is being curated. Third, experiments demonstrating the dbTF function are not available for many presumed dbTFs, and evidence based on homological proteins can sometimes be misleading, as for example the HLH proteins ID1-ID4 have lost their DNA binding capability and function instead as inhibitors of related dbTFs. Fourth, presence of even a functional DNA-binding domain does not always imply that the protein is a dbTF, as the GO definitions require evidence for the protein to have DNA regulatory activity besides it being able to bind DNA in a sequence-specific manner. Besides dbTFs there are many other proteins that bind to DNA with high binding affinities and have binding preferences for certain sequences over others, including the TATA binding protein, TBP, AT-hook proteins and even histones. Lastly, in some cases it is difficult to distinguish a dbTF from coTFs or GTFs, and occasionally, a protein is capable of more than one of these functions (16). In the course of this work, new guidelines have been developed to address those common pitfalls (Gaudet et al., 2020b, in preparation). In line with the view of many scientists working in this field (17), the term ‘DNA-binding transcription factor activity’ or a descendant term was applied only to those proteins that regulate transcription through the recognition of the genomic address of their target genes. This regulation is mediated by sequence-specific double-stranded DNA binding involving short cognate DNA motifs (18,19).

This article is focused on cataloguing sequence-specific DNA-binding transcription factors based on available evidence from experiments reported in literature, protein signatures, as well as by phylogenetic-based computational approaches. To create a compendium of human sequence-specific DNA-binding transcription factors, seven articles by experts in the field that list human dbTFs were selected (13,14,20–24). The 2,036 proteins cumulatively present in those lists were compared with existing GO associations, to yield an additional 61 candidate entries. Discrepancies were reviewed, which led to the generation of the present catalogue of 1,457 human dbTFs. Missing GO annotations were created, and incorrect annotations were removed. The human dbTF catalogue can be accessed via the GO browsers AmiGO2 (7) and QuickGO (25) or directly downloaded from https://www.ebi.ac.uk/QuickGO/targetset/dbTF and Supplementary Table S1A.

## Methods

### Resources used to create a list of potential human dbTFs

Potential human dbTFs were extracted from the supplementary files of five published articles (13,14,21–23) and two online resources, TFcheckpoint (20) and HumanTFDB (24), downloaded on 11 April 2020. The TFcheckpoint data was restricted to proteins for which literature potentially supporting their role as dbTF had been identified. Finally, the QuickGO browser (25) (https://www.ebi.ac.uk/QuickGO/) was used to download all human reviewed UniProt identifiers (IDs) annotated to either of the following three GO terms (or descendant terms): ‘DNA-binding transcription factor activity’ (GO:0003700), ‘transcription coregulator activity’ (GO:0003712), ‘general transcription initiation factor activity (GO:0140223)’, using the filters: Taxon - 9606, Gene Product - Protein, Reviewed, on 16 September 2019.

To create a dbTF comparison table (Supplementary Table S1), the resulting protein lists were aligned using the HUGO Gene Nomenclature Committee (HGNC) (26) approved gene symbol, gene name, and the UniProt ID (27). The majority of HGNC symbols and UniProt IDs were extracted using the UniProt Retrieve/ID mapping tool (28), although in some cases the HGNC multi-symbol checker (29) or Ensembl Biomart (30) were used.

### Assessment of dbTF function

The evidence supporting the assignment of the protein as a dbTF was reviewed. This often involved an extensive search of the published literature to identify robust experimental evidence of sequence-specific double-stranded DNA-binding, as well as evidence of regulation of transcription. GO annotation errors identified during the review were reported to the contributing group using GitHub (https://github.com/geneontology/go-annotation/projects/9) and corrected. In cases where the evidence for sequence-specific DNA binding was weak (Gaudet et al., 2020a, in preparation), additional support for dbTF activity was obtained from published High-Throughput Systematic Evolution of Ligands by Exponential Enrichment (HT-SELEX) data (31). This HT-SELEX data confirmed binding to specific DNA motifs for 540 human dbTFs. As this study uses recombinant proteins produced in *E. coli*, it provides strong evidence for direct DNA-binding, since very few complexes are expected to form between human dbTFs and endogenous *Escherichia coli* proteins (32–35).

To increase coverage of GO annotations, the PAINT (Phylogenetic Annotation and INference Tool), was employed to assign dbTF-associated annotations across species (11). Annotations were propagated only when there was convincing experimental support for at least one member of the PANTHER family or subfamily. Subfamilies were considered separately for large families or families with members with different functions, such as zinc finger-containing proteins. The annotations created using PAINT are associated with the IBA evidence code (the biological aspect of ancestor evidence used in manual assertion, IBA in Evidence and Conclusions Ontology (36)).

## Results

To create the catalogue of human dbTF based on the most current knowledge, a cumulative set of 2,097 putative human dbTFs was compiled from the resources described in the Methods (13,14,20–24) (Table 1, Supplementary Table S1). A variety of approaches were then taken to ensure that all human dbTFs were associated with an appropriate dbTF GO term.

**Table 1.**
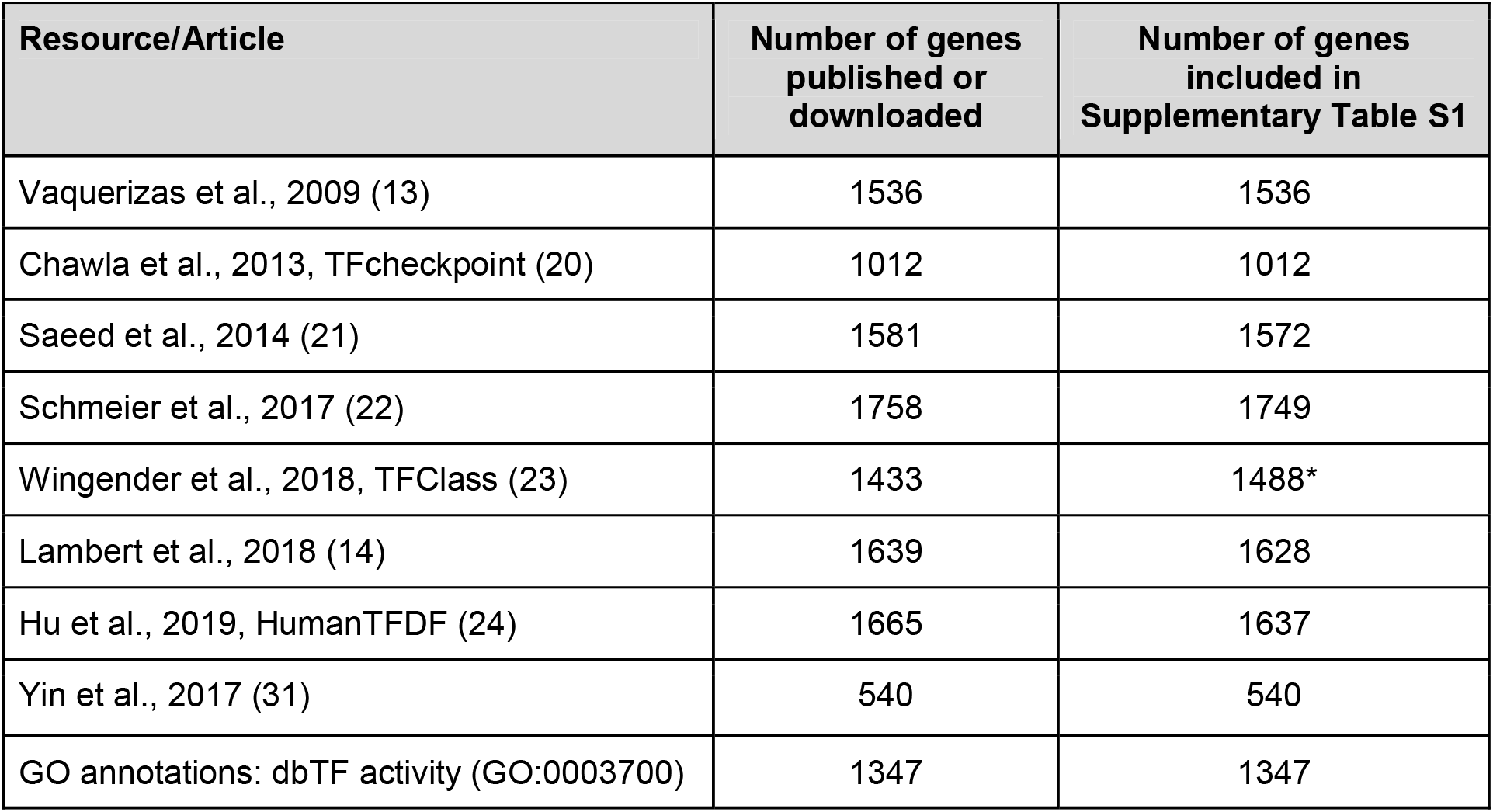
Summary of resources included in the dbTF comparison table. The number of dbTFs downloaded from each article or database (HumanTFDF, http://bioinfo.life.hust.edu.cn/HumanTFDB/, and TFcheckpoint, http://www.tfcheckpoint.org, downloaded on 11 April 2020, GO annotations https://www.ebi.ac.uk/QuickGO/ downloaded on 16 September 2019) is provided, as well as the number of genes included in the Supplementary Table S1. Mapping discrepancies were due to changes in genome builds and UniProt records. *Additional TFClass information was added to 55 genes in Sept 2020, following an update of this resource.

### Review of data supporting GO annotations

A manual review of all candidate human dbTFs associated with a dbTF activity GO term was undertaken to confirm the existing assignment as dbTF for each of these proteins. New annotation guidelines were formulated during the review of these annotations, guided in part by concrete examples of confirmed dbTFs, coTFs, and GTFs. These guidelines provided a strategy to enable curators to distinguish experimental evidence supporting the assignment to these molecular activity classes (Gaudet et al., 2020b, in preparation). Moreover, as HT-SELEX data included in the comparison table confirmed the ability of 540 human dbTFs to bind a valid discriminating DNA sequence-specificity (31), this information was occasionally used to inform the decision about whether a protein was likely to be a dbTF.

Over 3,000 GO annotations were reviewed; 583 human proteins were added to the ‘DNA-binding transcription factor activity’ GO group, while 256 human proteins were removed. All annotations were considered, i.e. manual annotations as well as automatic annotations, such as those based on UniProt keywords, InterPro domains, and annotations mapped from orthologous genes (9,11). The new GO transcription annotation guidelines were applied during the review of existing molecular function GO annotations describing transcription regulators and the revisions were implemented in the next GO annotation release. To complete this process, some non-human ortholog manual annotations were also revised, as these supported annotations that were associated with human orthologs. Requests for annotations to be changed were submitted to the relevant resource and tracked using GitHub (see methods section). As the GO Consortium has automatic annotation pipelines, the updated GO annotations associated with human proteins and InterPro records have been propagated to model organism proteins, thus improving these resources.

### Review of internally inconsistent GO annotations

Next to the RNA polymerases, GO defined three key molecular functions for proteins involved in transcription and its regulation: GTF, dbTF and coTF. Few proteins perform more than one of these functions and dbTFs rarely have catalytic activity, consequently, co-annotation to any combination of these terms was taken to indicate a possible misannotation.

#### Review of proteins annotated to dbTF and catalytic activity

The first review tested the hypothesis that while a substantial number of coTFs are enzymes, only a minority of dbTFs are also catalytically active enzymes. Following a download of all human protein annotations to either the GO:0003700, ‘DNA-binding transcription factor activity’ or GO:0003824 ‘catalytic activity’ and their descendant GO terms from QuickGO (25), a pivot table was used to identify 70 human proteins associated with a dbTF GO term as well as a catalytic activity term, leading to the review of over 450 annotations. Support for both activities was available for only 17 proteins, including CLOCK (37,38), several PRDM family members, and a few E2 ubiquitin-like protein conjugating ligases (including E4F1, EGR2, NFX1, ZBED1). The annotations associated with the remaining proteins were edited, with dbTF activity terms removed from 53 proteins, of which 11 should not have been annotated to either activity (Supplementary Table S1).

#### Review of proteins annotated to dbTF and coTF or GTF

Although dbTFs can also have coTF or GTF functions, this is not commonplace, therefore the second annotation review focused on human proteins associated with both a dbTF and a coTF or a GTF molecular function GO term. To identify these, a pivot table of human proteins annotated to one of the dbTF, coTF or GTF activity GO terms was generated. This identified 199 human proteins associated with the GO term ‘DNA-binding transcription factor activity’ that were also associated with either GO:0003712 ‘transcription coregulator activity’ or GO:0140223 ‘general transcription initiation factor activity’, or one of their descendants. As it was unlikely that the human proteome would include 200 dbTFs with two or more of these activities, a systematic review of the data supporting almost 1,400 GO annotations associated with these proteins was undertaken. In each case, the new GO annotation guidelines were applied (Gaudet et al., 2020b, in preparation). This led to the confirmed association of a dbTF activity term with 144 proteins and the removal of coTF or GTF terms from these proteins. Conversely, the dbTF term was removed from 55 proteins, the majority of which retained a coTF or GTF annotation or were annotated as dbTF inhibitors.

### Comparison of the seven sources of dbTF assignments

The comparison of seven lists of dbTFs demonstrates a considerable lack of consensus across these resources (Supplementary Table S1). Out of the 2,036 putative dbTFs, 818 human proteins were identified by all seven lists (40%), and 519 proteins were on six lists (25%), making up two-thirds of the initial set (Figure 1). A review of the experimental support for individual proteins or protein family members has confirmed that all but 44 of these 1,337 proteins are dbTFs. Many of the 44 excluded proteins have been assigned as coTFs, general chromatin structural proteins, or dominant negative inhibitors of DNA binding via heterodimer formation (Supplementary Table S1B). In contrast, a review of the literature describing 699 proteins described as dbTFs by one to five lists led to only 160 of these proteins being included in the dbTF catalogue (Supplementary Table S1A). This comparison highlights that proteins present in multiple lists are more likely to be dbTFs.

**Figure 1.**
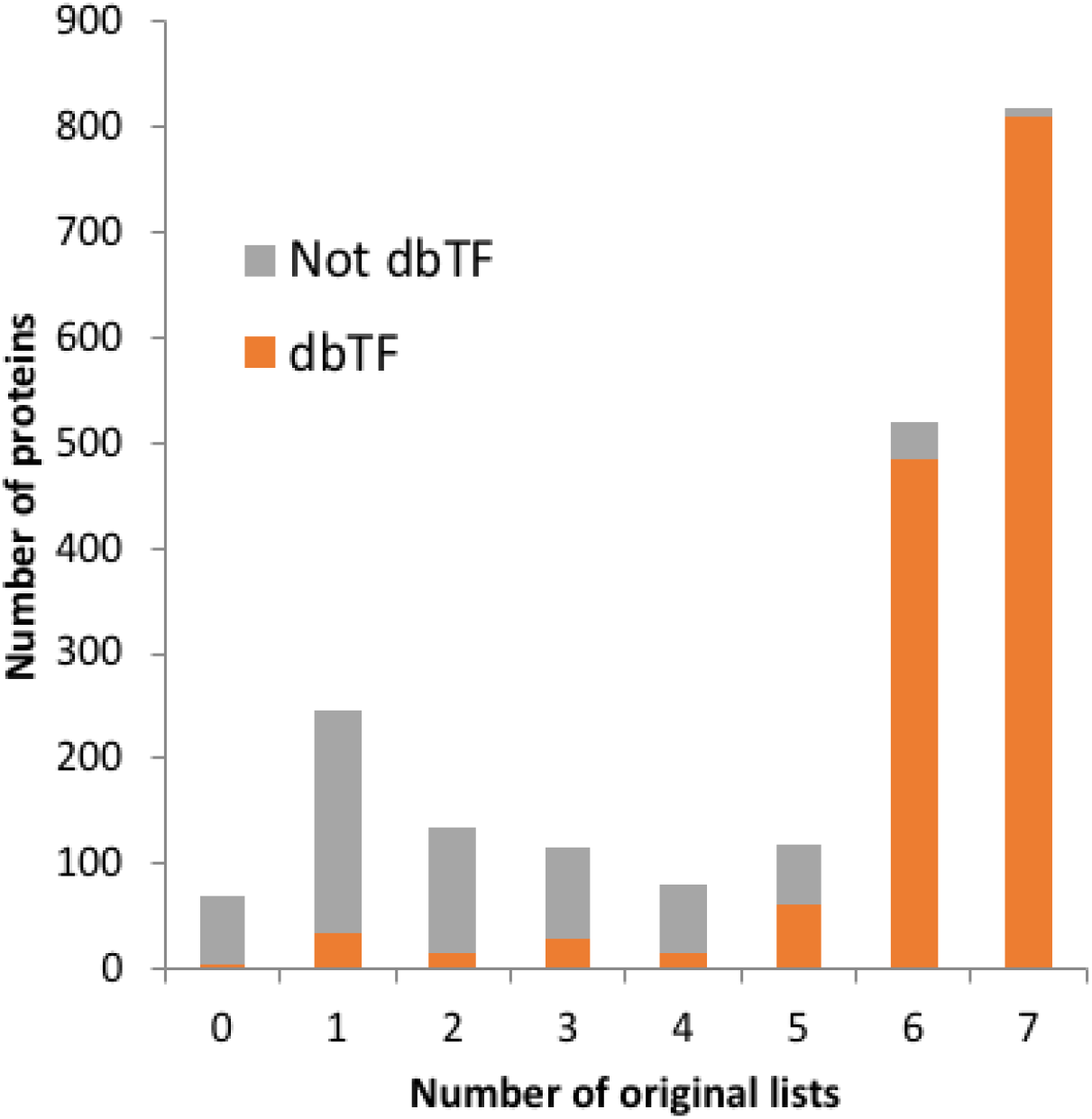
Representation of the number of proteins present in the compared lists. The 2097 proteins in the dbTF list comparison table (Supplementary Table S1) plotted as a function of the number of lists in which they are present (13,14,20–24).

The orange colour indicates the number of proteins that are included in the human dbTF catalogue (October 2020).

#### Review of proteins annotated to dbTF and included in five or fewer dbTF lists

The next approach was to manually review 169 proteins that were still associated with a dbTF GO term but were present in five or fewer of the dbTF lists. At this point, the literature supporting the association of a dbTF term with 41 proteins that were not present on any of the compared lists was also reviewed. Of these 210 proteins, 60 were confirmed to be dbTFs, the dbTF annotations associated with the remaining 150 proteins were removed. Three dbTFs were added to the human dbTF catalogue that had not been identified in any of the seven dbTF lists: the forkhead factors FOXL3 and FOXO3B, and the intracellular cleavage product of Junctophilin-2 (JPH2) (39).

This left 455 proteins that were not associated with any dbTF activity GO terms, but which were present on one to five of the dbTF lists. A review of curated knowledge provided by UniProt and InterPro (10,27) was conducted to determine if there was evidence to support the annotation of these 455 proteins as dbTFs. When this approach did not provide sufficient information, a literature review was undertaken. Ultimately, of those 455 proteins, 95 were included on the dbTF catalogue, although only 5 of these had experimental support. The remaining 90 were assigned as dbTFs based on InterPro protein signatures and/or by phylogenetic-based inference (11).

### Review of dbTF PANTHER protein families

The final step undertaken was to manually review PANTHER protein families and assign dbTF-relevant GO terms to proteins where there was evidence of conservation of the dbTF activity in that family. This was achieved using the GO Consortium PAINT tool (11). While this approach to curate PANTHER families has been established for over ten years, it had not been systematically applied to all dbTF families. For this project, we have curated 30 new dbTF Panther families and removed the dbTF annotations from 16 families. We have added dbTF annotations to 807 human proteins, for a total of 1,369 human proteins now having dbTF PAINT annotations, corresponding to 95% of the 1,440 dbTFs in the catalogue.

As some candidate dbTF PANTHER families had no dbTF activity GO annotations, an additional literature review was conducted and new, experimentally supported, GO annotations were created. Notably, PAINT annotations were also extended to non-human proteins resulting in the association of a dbTF activity GO term with 58,934 proteins, for 142 taxa. This review identified one dbTF, NFILZ, which was not present on any of the dbTF lists and had not been previously associated with a dbTF activity GO term.

### Examples of challenging protein families

This project has limited the assignment of a dbTF GO term to only sequence-specific double-stranded DNA-binding proteins that regulate transcription through the identification of the ‘genomic address’ of their target genes by binding to cognate short DNA motifs in the regulatory regions of these genes. In general, this strategy was relatively straightforward to apply. However, the presence of a DNA-binding domain does not always imply the protein is a dbTF, since DNA-metabolising enzymes also bind DNA, albeit not in a sequence-specific fashion. Therefore, direct *in vitro* data on DNA-binding specificities, such as HT-SELEX(31,40) provided a valuable additional source of experimental evidence to support some of the more challenging dbTF decisions. A specific DNA motif has been assigned to 1,007 of the human dbTFs (32). Notably, there are very few exceptions to the notion that sequence-specific DNA-binding implies that a protein is a dbTF. One interesting exception is the histone methyltransferase PRDM9, with 13 zinc fingers, that does bind DNA in a sequence-specific fashion but serves to mark meiotic recombination hotspots. Therefore, PRDM9 was not annotated as a dbTF (41) (Supplementary Table S1B). TERF1 provides another example of a protein inappropriately included in dbTF lists. This protein includes a homeobox-like domain and inhibits telomere elongation by sequence-specific binding of telomere ends (42).

The majority of challenging decisions involved excluding from the dbTF catalogue those proteins which contribute to the regulation of transcription through binding to single-stranded DNA (43), or structurally alter DNA conformation (44) (Supplementary Table S1B). In addition, proteins that bind short or unstructured, often AT-rich, sequences, such as ARID5A (45) and KDM5A (46), were not assigned a dbTF GO term. Many of these proteins function to increase the affinity of a true dbTF to the regulatory region of the genome rather than to provide the ‘genomic address’ and most are integral subunits of established transcription co-regulator complexes that remodel chromatin.

Another identified problem was contradictions in the literature. For example, Liefke et al., 2010 (46), stated that they could not reproduce the double-stranded DNA sequence-specific binding reported by Tu et al., 2008 (47) for the lysine demethylase KDM5A. Based on the report by Liefke et al., 2010, we chose to not annotate KDM5A as a dbTF. This type of inconsistency was addressed by considering whether or not the protein had sufficient experimental data to support the dbTF annotation.

All of the compared dbTF lists included zinc finger-containing proteins, which had led to the previous over-assignment of dbTF activity to this class of proteins (Supplementary Table S1B). As noted by Lambert et al., 2018 (14), these are the most challenging proteins to assign as dbTFs. While zinc fingers often recognize specific sequences of double-stranded DNA, some zinc fingers mediate protein interactions (48), in particular, interactions with small proteins such as ubiquitin and SUMO (49,50). For zinc finger-containing proteins that have not been experimentally investigated, this work considered the conservation of dbTF activity within the PANTHER subfamily. This approach led to the inclusion of 535 of 544 human krüppel C2H2 zinc finger proteins in the dbTFs catalogue, of which 348 harbour a KRAB domain (51). Of the nine human krüppel entries that were not included (ZFP64, ZFP91, ZNF335, ZNF451, ZNF488, ZNF507, ZNF513, ZNF653, ZNF827), eight belong to three PANTHER families with conflicting data and thus insufficient evidence to annotate or propagate dbTF activity. Furthermore, 72 C2H2 zinc finger proteins were excluded from the final dbTF list as there was evidence that these proteins act within coTF complexes or have small GTPase activating activity and no documented DNA binding activity.

## Discussion

Following a review of more than 3,000 GO annotations to some 2,000 proteins from several hundred articles, 1,457 proteins have been confirmed as dbTFs (Supplementary Table S1A), of which 1,414 are currently associated with a dbTF GO term. The list of potential dbTFs was then annotated using the PAINT curation tool, which provided annotations to homologous proteins based on their membership of orthology families or subfamilies. This review led to the removal of the dbTF activity term from 256 proteins and a new assignment of dbTF activity to an additional 583 proteins.

The creation of a list of human dbTFs has been undertaken several times over the last decade involving many different groups of researchers (13,14,20–24,52). The difficulty of this task is highlighted by the limited number (65%) of sequence-specific DNA-binding transcription factors that are in the intersection of at least six of the seven lists we compared. Notably, ten proteins present in all seven lists are not considered by this study to be dbTFs. In addition, there were almost 600 proteins, present on at least one of these lists, for which there was no evidence of their role as a dbTF (Supplementary Table S1B). These discrepancies stem from a number of factors, including the weight the authors give to the available supporting evidence, the definition of transcription factors that was applied, and the criteria used to organise proteins in orthology subfamilies. Furthermore, some of the errors in the dbTF lists were likely to be due to the propagation of misannotations in the previous GO annotation files. The work presented here has addressed this problem by removing incorrect dbTF activity assignments.

Here we report a catalogue of human dbTFs that has been evaluated against rigorous annotation criteria that allows integration in GO. Despite the care we took, it is likely that this list harbours some false positives that will need to be removed in the future. Similarly, there are likely to be false negatives, namely true dbTFs which have yet to be identified and/or validated. However, as only four dbTFs were identified that had not been included on any of the compared dbTF lists, the number of false negatives is likely to be small. The present systematic review has substantially improved the public GO annotation files by applying a uniform approach to reviewing the existing knowledge of dbTFs and the supporting experimental data. This was achieved by consistently applying new annotation guidelines to a comprehensive list of candidate dbTFs (representing around 10% of the human protein coding genes) and concomitantly refactoring the transcription-relevant domains of GO. The GO curation effort had previously been impaired by the excessive number of GO terms describing the functions that contribute to transcription. In the course of this work, the molecular function ontology describing transcription regulators has been substantially revised, with around 90% of these GO terms either obsoleted or merged (Gaudet et al., 2020a, in preparation). In addition, the term definitions have been improved and new GO annotation guidelines have been created to promote appropriate annotation of dbTFs, coTFs, and GTFs (Gaudet et al., 2020b, in preparation). This may be the first time that an integrated approach for transcription factor activities, alongside adaptation of the Gene Ontology itself, was undertaken for the gene regulation knowledge commons.

As the prime regulators of a plethora of biological processes, dbTFs are of fundamental importance. The availability of a comprehensive list of dbTFs is an essential step in reconstructing the core layers of gene regulatory networks starting from the analysis of genuine dbTF binding sites in gene regulatory regions. In addition, interpretation of the results of high-throughput methodologies strongly depends on high-quality gene annotations. In particular, a clear distinction between dbTFs and coTFs will facilitate precise identification of causal regulatory variants, e.g. when using high-throughput data from chromatin immunoprecipitation or open chromatin assays. Consequently, we expect that the reviewed dbTF list presented here will benefit the systems biology and biomedical research communities as well as facilitate fundamental proteomic, transcriptomic, and genomic research.

The catalogue of human dbTFs presented here, along with the GO term revisions, transcription annotation guidelines, and review of annotations, will aid future curation efforts of dbTFs across all species and ensure that the Gene Ontology accurately describes our current understanding of transcription and the regulation of this process. The dbTF catalogue includes 1,414 dbTFs that are associated with a dbTF activity GO term based on published experimental evidence or membership in a well characterised dbTF family. However, the proposed list of dbTFs demonstrates that further research in this area is still required. Only 515 human dbTFs are assigned a dbTF activity GO term based on experimental evidence. There are still around 900 human dbTF with no direct experimental verification of their role as dbTFs. We, therefore, call on the transcription research community to interrogate GO and to target the under-investigated candidate dbTFs to provide biochemical verification of the role of these proteins as dbTFs. In addition, articles describing negative data, which excludes the dbTF activity of a protein, are invaluable for clarification of dbTF assignment, but often difficult to find. The GO Consortium would welcome information from interested researchers about existing, but not currently curated, high-quality data that provide experimental support for these under-annotated proteins, so as to improve this resource.

### Availability of the dbTF catalogue

The human dbTF catalogue is available as Supplementary Table S1A and can be downloaded from the new webpage (https://www.ebi.ac.uk/QuickGO/targetset/dbTF). In addition, all GO dbTF activity annotations are freely available to download or to browse using the GO browsers AmiGO2 (7) (http://amigo.geneontology.org/amigo/) filtering on GO term ‘GO:0003700’, Ontology (aspect) ‘F’, organism ‘Homo sapiens’ or QuickGO (25) (https://www.ebi.ac.uk/QuickGO/) filtering on GO terms ‘GO:0003700’, taxon ‘human’, gene products ‘proteins Reviewed (Swiss-Prot)’.

## List of Abbreviations

coTF: transcription coregulator
dbTF: DNA-binding transcription factors that bind to a specific-sequence (or motif) in double-stranded DNA to provide a genomic address.
GO: Gene Ontology
GTF: general (or basal) transcription factor

## Acknowledgements

We apologise to those researchers whose relevant work we did not cite. Thanks to Dustin Ebert for his help with PAINT and PANTHER and thanks to many biocurators who promptly revised GO annotations when revisions were requested, including Hsin Yu Chang, Sara Chuguransky, Harold Drabkin, Li Ni, Karen Christie, David Hill, Penelope Garmiri, Michele Magrane, Sylvain Poux, Cristina Cassals, Lionel Breuza, Ghislaine Argoud-Puy, Thomas Hayman, Helen Attril.

This article is based upon work from COST Action CA15205, supported by COST (European Cooperation in Science and Technology). RCL was supported by the British Heart Foundation (RG/13/5/30112), Alzheimer’s Research UK (ARUK-NAS2017A-1), and the National Institute for Health Research University College London Hospitals Biomedical Research Centre. IVK was supported by Russian Science Foundation grant 20-74-10075. OF was supported by the BC Children’s Hospital Foundation and Research Institute. MLA was supported by the Research Council of Norway, project number 247727/O70. The GO Consortium and PG are funded by the National Human Genome Research Institute (US National Institutes of Health), grant number HG002273.

## Supplementary Table S1 legend

**Supplementary Table S1. A comparison of dbTF lists.** A comparison of dbTF molecular function GO annotations with three resources and five articles providing lists of human dbTFs. Table S1A. Human dbTF catalogue: identifies the genes this review has assigned as dbTFs https://www.ebi.ac.uk/QuickGO/targetset/dbTF; Table S1B. Reviewed genes not dbTFs: identifies the genes this review does not consider likely to be dbTFs. Column headings for both Tables: HGNC approved gene symbol; HGNC approved gene name; Pseudogene: proteins that have been assigned pseudogene status either by HGNC or UniProt; UniProt ID: UniProt identifier; dbTF GO annotation 16Sept2019 (1347 genes): 1347 genes associated with a dbTF activity GO term downloaded on 16 September 2019; Vaquerizas, 2009 (1536 genes) with dbTFs categories (13); TFCheckPoint, Chawla, 2013 (1012 genes) downloaded on 11 April 2020 from TFCheckPoint (20); Saeed, 2014 (1572 genes) (21); Schmeier, 2017 (1750 genes) (22); TFClass, Wingender, 2018 (1488 genes), TFClass IDs are based on the characteristics of their DNA-binding domains with four general levels (superclass, class, family, subfamily) and two levels of instantiation (genus and molecular species), subfamily and factor species are not provided for all proteins (23); Lambert, 2018 (1630 genes) (14); HumanTFDB, Hu, 2019 (1637 genes) downloaded on 11 April 2020 from HumanTFDB (24); Number of dbTF lists: the total number of dbTF list the gene is included in (columns J-P); HT-SELEX, Yin, 2017 (540 genes) genes identified by HT-SELEX data as binding double-stranded DNA(31); PANTHER family ID, subfamily ID and family name assigned by PANTHER, www.pantherdb.org (53).

All dbTF activity GO annotations are freely available to download or browse using the GO browsers AmiGO2 (7) (http://amigo.geneontology.org/amigo/) filtering on GO term ‘GO:0003700’, Ontology (aspect) ‘F’, organism ‘Homo sapiens’ or QuickGO (25) (https://www.ebi.ac.uk/QuickGO/) filtering on GO terms ‘GO:0003700’, taxon ‘human’, gene products ‘proteins Reviewed (Swiss-Prot)’. Furthermore, the catalogue of human dbTFs presented here is available for downloading from the QuickGO browser (https://www.ebi.ac.uk/QuickGO/targetset/dbTF).

While there are small differences between the lists of dbTFs downloaded from the different resources, Supplementary Table S1A contains the full list of dbTFs. The differences between these dbTF lists arise from the lack of GO annotations or UniProt IDs. There is no experimental data for 26 proteins or their family members. The dbTF catalogue includes 28 pseudogenes as these may code functional proteins in some individuals (54). As some of these pseudogenes do not have a UniProt ID they are not present in the QuickGO or AmiGO2 dbTF lists. In addition, two proteins (LITAF and NDN) are associated with a dbTF activity GO term despite having no obvious DNA-binding domain. In these cases, the dbTF GO annotations were not contested, but the data were not considered sufficiently robust for these proteins to be included in the current dbTF catalogue.

